# Mapping of genomic regions linked to stemphylium blight (*Stemphylium botryosum* Wallr.) resistance in lentil using linkage mapping and marker-trait association analysis

**DOI:** 10.1101/2022.08.22.504834

**Authors:** Stanley Adobor, Tadesse S. Gela, Sabine Banniza, Albert Vandenberg

## Abstract

Stemphylium blight caused by *Stemphylium botryosum*, is a foliar disease of lentil. It affects the productivity and milling quality of lentil crops, mainly in South Asia and Canada. Development of stemphylium blight resistant cultivars by introgression of resistance alleles from crop wild relatives of lentil, such as *Lens ervoides*, is one strategy of disease control. The objective of this study was to identify genomic regions associated with stemphylium blight resistance by combining linkage mapping and marker-trait association analysis. A total of 182 genotypes of a lentil advanced backcross population (LABC-01) developed from the backcross of the interspecific *L. culinaris × L. ervoides* line LR-59-81 (donor) and cultivar CDC Redberry (recurrent) and 101 diverse lentil accessions selected by stratified random sampling from a lentil diversity panel were genotyped and evaluated for stemphylium blight reactions. Quantitative trait locus (QTL) analysis identified four loci contributing to stemphylium blight resistance on lentil chromosomes 2, 4 and 5. Marker trait association analysis detected five significant single nucleotide polymorphism (SNP) markers associated with stemphylium blight resistance within QTLs regions and seven SNP markers outside the QTLs regions on chromosomes 1, 2, 3, 5, and 7. The markers associated with stemphylium blight resistance may be useful for marker-assisted selection of resistant cultivars after validation.

## 1 Introduction

Cultivated lentil (*Lens culinaris* ssp *culinaris* Medik.) is a diploid (2n = 2x = 14), self-pollinating annual pulse crop with a genome size of approximately 4 Gbp (Arumuganathan and Earle, 1991). It is grown in 70 countries with a total annual production of 6.54 Mt from an estimated 5.0 M ha with an average yield of 1.31 t ha^−1^ (FAO, 2020). It is an important source of protein, carbohydrates, and micronutrients in human diets and is a widely consumed pulse crop, with a rate of annual consumption growing much faster than that of human population growth (Khazaei et al., 2019). Production of lentil is, however, constrained by biotic and abiotic stresses. Among the biotic factors, fungal diseases are the most important constraint to lentil productivity.

Stemphylium blight, commonly associated with the fungal pathogen *Stemphylium botryosum* Wallr., threatens lentil productivity in the major growing regions, including Canada, India, Australia, Turkey, Nepal, and the USA (Erskine and Sarker, 1996; Bayaa and Erskine, 1998; Morrall et al., 2004). It is a foliar disease responsible for large-scale rapid defoliation of lentil plants, resulting in a loss of yield in conducive environments. Stemphylium blight epidemics are favoured by warm temperatures between 25 C and 30 C combined with conditions of more than 85% humidity for 48 h (Mwakutuya and Banniza, 2006). Yield losses of up to 62% have been reported in India, Nepal, and Bangladesh (Taylor et al., 2007). *S. botryosum* was frequently detected on lentil seeds tested in commercial seed testing laboratories in Canada (Morrall et al., 2006). In addition, late-season disease infection has been reported to reduce milling quality (Subedi et al., 2021). Improving resistance to stemphylium blight in lentil crops is therefore crucial for stable food production.

Crop wild relatives are an important source of novel genes lost during the domestication process (Tanksley and McCouch, 1997; Haussmann et al., 2004; Hajjar and Hodgkin, 2007; Maxted et al., 2008; Ford-Lloyd et al., 2011; Warschefsky et al., 2014). In the four gene pools of the *Lens* genus (Wong et al., 2015), wide crosses of lentil cv. Eston with *L. ervoides* (Brign) Grande accessions belonging to the tertiary gene pool were made to transfer desirable genes for disease resistance and to increase the genetic variability in cultivated lentil (Fiala et al., 2009; Tullu et al., 2013). Inheritance of resistance to anthracnose (Fiala et al., 2009; Tullu et al., 2013; Gela et al., 2021a) and stemphylium blight (Adobor et al., 2020) have been studied in *L. ervoides* interspecific recombinant inbred line (RIL) populations.

Identification of quantitative trait loci (QTLs) or genes controlling stemphylium blight resistance and linked molecular markers is a primary step in efforts to improve lentil cultivars by introgression of beneficial QTLs from wild resistance sources into cultivated backgrounds. However, characterization of the genetic control of stemphylium blight resistance using interspecific RIL populations has been hampered by segregation distortion. Utilization of interspecifics in breeding programs has also been limited by linkage drag, making selection practically slow. Tullu et al. (2013) and Chen (2018) reported undesired traits such as seed dormancy, poor emergence, extremely small seed size, and seed shattering in *L. ervoides* interspecific RILs. One of the approaches to minimizing the transfer of undesirable traits is the use of advanced backcrossing combined with QTL analysis for simultaneous molecular breeding and QTL mapping (Tanksley and Nelson, 1996). It relies on the partial isolation of wild QTLs by repeated backcrosses to an adapted parent to reduce much of the wild genome in the cultivated background. This approach has been applied to many crops (Frary et al., 2004; Silvana and Tanksley, 2005; von Korff et al., 2005; Yu et al., 2005; Narasimhamoorthy et al., 2006; Naz et al., 2008; Schmalenbach et al., 2008; Bauer et al., 2009) including lentil (Gela et al., 2021b).

The development of molecular markers in the 1980s (Tanksley and Rick, 1980; Paterson et al., 1988) has made the identification of QTLs and mapping of complex traits to specific chromosome region(s) feasible by linkage mapping using bi-parental mapping populations (Collard et al., 2005) or by linkage disequilibrium (LD) mapping using natural populations (Brachi et al., 2010). Linkage mapping is highly dependent on the genetic diversity of the two parents and recombination in the mapping population (Borevitz and Nordborg, 2003; Collard and Mackill, 2008; Zhu et al., 2008; Corwin and Kliebenstein, 2017). LD mapping takes advantage of historic recombination in natural populations and evaluates numerous alleles to detect QTLs at a high resolution (Nordborg and Tavaré, 2002; Zhu et al., 2008; Brachi et al., 2010; Corwin and Kliebenstein, 2017). Both approaches have been combined to precisely map QTLs in maize and lentil (Mammadov et al., 2015; Chen et al., 2016; Wu et al., 2020; Gela et al., 2021c). With the advances in sequencing and the availability of the reference genome of lentil cultivar CDC Redberry (Ramsay et al., 2021), single nucleotide polymorphism (SNP) discovery and QTL mapping can be performed, and QTLs can be more precisely localized on the genetic and physical sequence maps.

In this study, linkage mapping was applied to identify genomic regions associated with stemphylium blight resistance in an advanced backcross population, LABC-01 (Gela et al., 2021b), that was evaluated in three environments. Next, genome-wide association analysis was performed to identify SNP markers linked to stemphylium blight resistance within and outside of the QTL-regions. The objectives of this study were to identify genomic regions underlying stemphylium blight resistance in *L. ervoides* interspecifics and to identify linked molecular markers that may be used by breeders to develop resistant cultivars.

## 2 Materials and Methods

### 2.1 Plant material

Genetic resistance to *S. botryosum* was evaluated using an advanced backcross population (LABC-01) and a genome-wide marker trait association panel. LABC-01 was derived from a cross between interspecific RIL LR-59-81 (*L. culinaris* ‘Eston’ × *L. ervoides* L01-827A) and ‘CDC Redberry’ (Gela et al., 2021b) and was comprised of a set of 182 individual BC_2_F_3:4_ genotypes. The donor line LR-59-81 was previously identified to be resistant to *Collectotrichum lentis* Damm races 0 and 1 (Fiala et al., 2009; Gela et al., 2020) and stemphylium blight. The recurrent parent, CDC Redberry, was one of the first small red lentil cultivars released by the Crop Development Centre (CDC), University of Saskatchewan. CDC Redberry has resistance to both ascochyta blight and anthracnose race 1, combined with excellent lodging tolerance and high yield (Vandenberg et al., 2006). CDC Redberry was also moderately resistant to stemphylium blight (unpublished data). The lentil diversity panel for marker trait association analysis consisted of 101 *L. culinaris* genotypes representing a stratified random sample from a diversity panel of 324 accessions assembled previously (Haile et al., 2020; Gela et al., 2021c; full list available at http://knowpulse.usask.ca/Lentil-Diversity-Panel). The collection was originally obtained from the gene banks of the International Center for Agricultural Research in the Dry Areas (ICARDA), the United States Department of Agriculture (USDA), Plant Gene Resources of Canada (PGRC), and cultivars developed at the CDC.

### 2.2 Disease phenotyping

The LABC-01 population was evaluated in three environments: the greenhouse in 2018, a growth chamber (GR178, Conviron, Winnipeg, MB, Canada) in 2019 in the College of Agriculture and Bioresources phytotron facility of the University of Saskatchewan, Canada, and a field experiment at the Seed Farm of the Department of Plant Sciences in 2019. All three experiments were performed in a randomized complete block design with three replicates. The lentil diversity panel was evaluated in a growth chamber in 2019 with a randomized complete block design in four replicates. Cultivars Eston (Slinkard, 1981) and CDC Glamis (Vandenberg et al., 2002) were included in all experiments as moderately resistant and susceptible checks, respectively. Plant growth conditions and maintenance for experiments in the greenhouse and growth chamber were carried out as described by Adobor et al. (2020) and Gela et al. (2021b). Briefly, six seeds from each LABC-01 population line, parents, and checks were planted in 10-cm plastic pots filled with SUNSHINE MIX #4 plant growth medium (Sun Gro Horticulture, Seba Beach, AB, Canada). Plants were thinned to four plants per replicate pot two weeks after emergence and fertilized once a week with 3 g L^−1^ of soluble N:P:K (20:20:20) PlantProd® fertilizer (Nu-Gro Inc., Brantford, ON, Canada). A culture stock of *S. botryosum* isolate SB19, obtained from the Pulse Crop Pathology program of the CDC, was used for mass spore production. Plants were spray-inoculated at the pre-flowering stage using approximately 3 mL of *S. botryosum* isolate SB19 conidial suspension per plant at a concentration of 1 × 10^5^ conidia mL^−1^ using an air brush (Badger Airbrush, model TC 20, USA). Two drops of Tween^®^ 20 (Sigma, Saint Louis, MO, USA) were mixed with conidial suspensions to reduce the surface tension of water and promote plant tissue contact. Plants were placed in an incubation chamber. Two humidifiers (Vicks Fabrique Paz Canada, Inc., Milton, ON, Canada) were placed in the incubation chamber to maintain a high relative humidity conducive for infection and disease development.

Stemphylium blight severity was assessed visually at 7 days post inoculation using a semi-quantitative rating scale (0-10) where 0, healthy plants; 1, few tiny lesions; 2, a few chlorotic lesions; 3, expanding lesions on leaves, onset of leaf drop; 4, 1/5th of nodes affected by lesions and leaf drop; 5, 2/5th of nodes affected; 6, 3/5th of nodes affected; 7, 4/5th of nodes affected; 8, all leaves dried up; 9, all leaves dried up but stem green; and 10, plant completely dead. Disease severity was assessed for each genotype on each of the four individual plants within the experimental unit (pot).

In the field, 12 seeds of each BC_2_F_3:4_ LABC-01 line were planted in the middle row between the border rows of the susceptible check CDC Glamis in a three-row micro plot (1 m x 0.6 m) seeding arrangement. Field management followed the standard agricultural practices for field grown lentil. For disease inoculation, ground faba bean seeds infected with SB19 spores were spread in the plots (~20g per plot) prior to spray inoculation to ensure enough inoculum was available for infection and disease development. At the pre-flowering stage, a knapsack sprayer was used to apply approximately 400 mL conidial suspension of SB19 at 1 × 10^5^ conidia mL^−1^ concentration per plot to further increase disease pressure. Plots were covered after inoculation with perforated green polyethylene low horticultural-tunnel covers (Dubois Agrinovation, Saint-Rémi, Quebec, Canada) to create a conducive microclimate for pathogen infection. When the susceptible check CDC Glamis was sufficiently diseased (61-70% severity), stemphylium blight severity was recorded from five randomly selected plants in the middle row using a quantitative rating scale ranging from 0 to 10 with 10% increments of disease severity (DS), where 0 = 0% DS, 1 = 1-10% DS up to 10 = 91-100% DS. Data were converted to percentage disease severity using the class midpoints for data analysis.

### 2.3 Disease severity and data analysis

All analyses of disease severity data were performed using SAS software version 9.4 (SAS Institute, Cary, North Carolina, USA). Disease severity data collected from the greenhouse, growth chamber and field (hereafter referred to as the environment) were analyzed separately. The data were checked for normality of residuals and homogeneity of variance. The REPEATED/GROUP statement was used to model heterogeneous variance where Levene’s test for homogeneity was significant. Genotype was modelled as a fixed effect and block was considered random factor using the PROC MIXED procedure. The LSMEANS statement was used to estimate least squares means for QTL analysis. The PROC VARCOMP procedure was used to estimate the variance components of the severity data. Broad sense heritability (H^2^) was estimated as 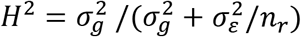 for each environment and as 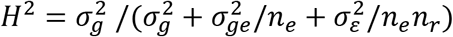 for combined greenhouse and growth chamber disease severity scores (Knapp et al. 1985), where 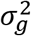 is the genotype variance, 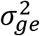 is the genotype by environment interaction variance, 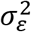 is the error variance, ne is the number of environments, and nr is the number of replications.

### 2.4 Whole-exome sequencing of LABC-01 population and Lentil diversity panel

Genomic DNA was extracted from freeze-dried leaves collected from single plants of the 182 BC_2_F_3:4_ genotypes and the parents, using the DNeasy Plant Mini Kit (Qiagen, Hilden, Germany) following the manufacturer’s protocol. Sequence library, read mapping, and SNP genotyping of the LABC-01 population and lentil diversity panel were performed using a custom lentil exome capture assay as described by Ogutcen et al. (2018). SNP calling was performed by mapping sequenced data to the *L. culinaris* cv. CDC Redberry genome assembly v2.0 (Ramsay et al., 2021). The SNP data for the 101 lentil accessions from the diversity panel were obtained from the KnowPulse website using the vcf bulk export tool (Sanderson et al., 2019; https://knowpulse.usask.ca/study/2675314).

### 2.5 Genetic map construction and QTL analysis

The linkage map of the LABC-01 population was constructed with 877 SNP markers for 182 BC_2_F_3:4_ individuals using the QTL IciMapping software V4.2 (Meng et al., 2015). A logarithm of odds (LOD) threshold of 3.0 was used to group SNP markers into linkage groups (LG). SNP markers within a LG were ordered using a recombination counting and ordering algorithm (RECORD). The marker order was refined by rippling with a number of recombination events (COUNT) algorithm with a window of five markers. Map distance in centiMorgan (cM) was calculated using the Kosambi mapping function.

QTL analyses were performed separately for each environment using mean severity data. The IM-ADD and ICIM-ADD mapping functions were used to implement the interval mapping and the inclusive composite interval mapping methods for QTL detection in the QTL IciMapping software. The LOD threshold for declaring significant QTL was estimated by running 1,000 permutations with a type I error at *P* ≤ 0.05. The QTL were named following the nomenclature of McCouch et al. (1997): denoted as q + trait name + chromosome + number of QTL on the chromosome, such as “*qSB-1-1*”, where “*qSB*” indicates the QTL for stemphylium blight and “*1*-*1*” indicates the first QTL on chromosome 1.

### 2.6 Marker-trait association analysis

SNP marker data for 101 lentil accessions were filtered using VcfTools v.0.1.15 with the following parameters: bi-allelic SNPs, minimum read depth of 3; minor allele frequency (MAF) ≥ 5%; and maximum missing frequency < 20% (Danecek et al., 2011). The remaining 461,411 SNP markers were further pruned using linkage disequilibrium (LD) values of *r^2^* > 0.8 between SNP markers assessed with PLINK 1.9 software (Purcell et al., 2007). A total of 143,971 SNPs remained after pruning. Flanking markers for four QTLs detected in the LABC-01 population (*qSB-2-1, qSB-2-2, qSB-4-1, qSB-5-1*) were obtained based on a 1.5-LOD confidence interval and used to delimit the physical QTL regions on the CDC Redberry reference genome v.2.0. A total of 2,416 SNP markers were identified in the QTL regions and used for analysis.

All marker-trait association analyses were performed using the R package GAPIT (Genome Association and Prediction Integrated Tool) version 3.0 (Wang and Zhang, 2018). The population structure was evaluated by Discriminant Analysis of Principal Components (DAPC) (Jombart et al., 2010) using the *adegenet* package (Jombart, 2008) for R software. The kinship matrix (K) was calculated using the algorithm implemented in GAPIT3 (VanRaden, 2008). Genome-wide and QTL-region (QTLs from the LABC-01 population) marker trait association analysis was performed using fixed and random model Circulating Probability Unification (FarmCPU) (Liu et al., 2016). The FarmCPU model was fitted using the 2,416 SNPs for QTL-regions and the 143,971 SNPs for GWAS and incorporating the kinship matrix (K) as a random effect and the first two principal components (PC) as fixed effect covariates to control spurious association. Significant SNP markers were identified based on FDR-adjusted *P* ≤ 0.05 (Benjamini and Hochberg, 1995). LD between SNP markers was calculated using PLINK 1.9. All R packages were run on R statistical software v. 4.1.2 (R Core Team, 2021).

## 3 Results

### 3.1 Stemphylium blight severity in LABC-01 population and lentil diversity panel

The LABC-01 population was screened under three different environments to evaluate variations in stemphylium blight severity. The interspecific donor line LR-59-81 is resistant to stemphylium blight, and the recurrent parent CDC Redberry is moderately resistant. The mean severity rating for LR-59-81 and CDC Redberry in the greenhouse were 2.17 and 4.17, and in the growth chamber, 2.33 and 5.17. In the field, the mean disease severity was 34.87% for LR-59-81 and 40.25% for CDC Redberry. The mean severity rating of BC_2_F_3:4_ individuals in the greenhouse and growth chamber ranged from 1.33 to 6.00 and 1.83 to 5.83. Severity in the field ranged from 25.30% to 63.43%. Significant genotype effects on disease severity were observed in the greenhouse (F-value = 13.62, *P* = 0.0001), growth chamber (F-value = 56.99, *P* = 0.0001), and in the field (F-value = 1.63, *P* = 0.0001). No resistant transgressive segregants were observed among the LABC-01 genotypes based on severity data from all three environments. The distribution of disease severity in the 182 BC_2_F_3:4_ individuals in all three environments showed a unimodal distribution pattern of segregation (Figure 1A - 1C), suggesting quantitative inheritance of stemphylium blight resistance in the LABC-01 population. The variance components for genotype, environment, and genotype by environment interaction effects on stemphylium blight severity were significant. The estimated broad sense heritability for each environment was high ranging from 40.1% in the field to 62.3% in the greenhouse, whereas the heritability for the combined severity data from the greenhouse and growth chamber was 16.1% (Table 1).

**Figure 1.**
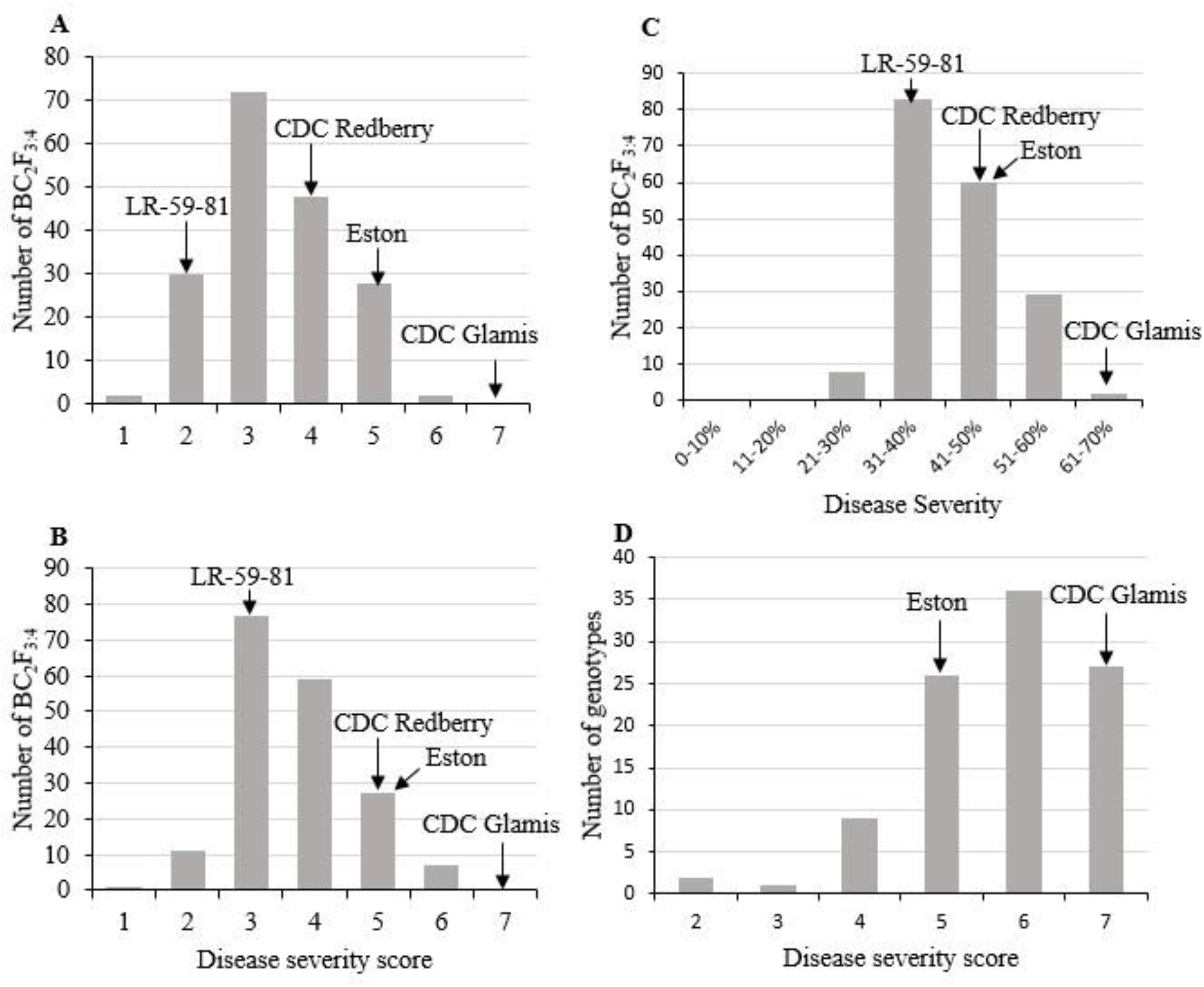
Frequency distribution of stemphylium blight severity in advanced lentil backcross population LABC-01 (n =182, BC_2_F_3:4_) evaluated in (A) greenhouse in 2018, (B) growth chamber in 2019 and (C) field in 2019, and (D) 101 accessions of a lentil diversity panel screened in the growth chamber.

**Table 1.** Analysis of variance components for stemphylium blight severity of 182 BC_2_F_3:4_ (CDC Redberry x LR-59-81) population evaluated under greenhouse, growth chamber and field conditions and 101 diversity panel evaluated in the growth chamber.

| Environment | Mean | Range | H<sup>2</sup> (%) | $\sigma_g^2$ | $\sigma_e^2$ | $\sigma_{ge}^2$ | $\sigma_\varepsilon^2$ | CV (%) |
| --- | --- | --- | --- | --- | --- | --- | --- | --- |
| **LABC-01** |  |  |  |  |  |  |  |  |
| Greenhouse-2018 | 3.3 | 1.33 - 6.00 | 62.3 | 0.522 * |  |  | 0.948 | 29.27 |
| Growth chamber-2019 | 3.64 | 1.83 - 5.83 | 46.7 | 0.296* |  |  | 1.014 | 27.64 |
| Field-2019 | 43.15 | 25.07 - 63.33 | 40.7 | 21.842* |  |  | 95.483 | 22.64 |
| Combined GH-GC<sup>a</sup> | 3.47 | 2.00 - 5.42 | 16.1 | 0.064 | 3.76 x 10<sup>-10</sup> | 0.344* | 0.981 | 28.51 |
| **Diversity panel** |  |  |  |  |  |  |  |  |
| Growth chamber-2019 | 5.4 | 2.63-7.00 | 65.4 | 0.561* |  |  | 1.187 | 19.9 |

H<sup>2</sup>; broad sense heritability, $\sigma_g^2$ ; genotypic variance, $\sigma_e^2$ ; environmental variance, $\sigma_{ge}^2$ ; genotype by environment interaction variance, $\sigma_\varepsilon^2$ ; error variance; CV%; coefficient of variance, <sup>a</sup> combine disease score data of greenhouse and growth chamber, \*; significant at $P \leq 0.05$ .

Stemphylium blight severity among the 101 diverse lentil accessions screened in the growth chamber ranged from 2.63 to 7.00 with a mean of 5.4 (Figure 1D). Variation in stemphylium blight severity due to genotypes was significant (F-value = 2.34, *P* = 0.0001).

### 3.2 Genetic map construction and QTL analysis

Whole exome sequencing of 182 LABC-01 individuals yielded 5,017 SNP markers polymorphic between the parents. The SNP markers were further filtered to 877 for significant segregation distortion and missing data. Of the 877 SNP markers, 727 (82.9%) were located within genes on the CDC Redberry reference genome v2.0.

The proportion of markers in the 182 BC_2_F_3:4_ individuals homozygous for the recurrent parent genotype was 84.9%, whereas 9.5% were homozygous for the donor genotype and 5.6% were heterozygotes. Using all 877 markers, the genetic map was estimated with QTL IciMapping software, which is more tolerant of distorted markers. The 877 markers were grouped into 7 linkage groups (LG) corresponding to the haploid chromosome number of *L. culinaris* using a LOD score ≥ 3 (Figure 2; Supplementary Table 1). The markers covered a total map distance of 2,566.6 cM with an average marker-interval of 3 cM. The total map length for each LG ranged from 66.1 cM (LG7) to 652.2 cM (LG4). Marker coverage ranged from 8 on LG7 to 326 on LG2. The average inter-marker gap ranged from 1.6 cM on LG2 (high marker density) to 9.4 cM on LG7 (low marker density).

**Figure 2.**
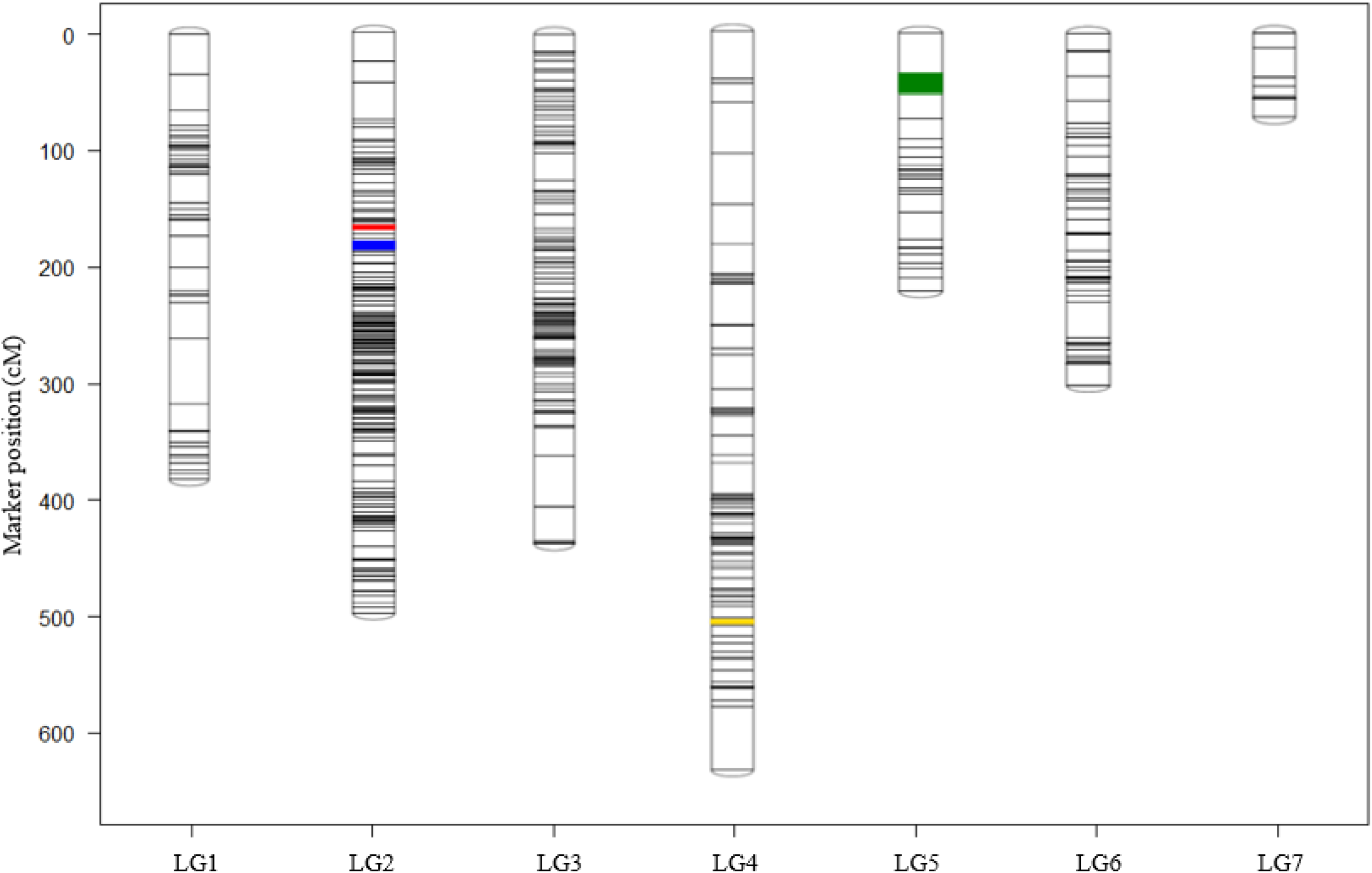
Linkage map based on 877 SNP markers shows QTL controlling stemphylium blight resistance in advance lentil backcross population LABC-01 (n =182, BC_2_F_3:4_). The colored regions represent 1.5-LOD confidence intervals of QTLs detected using stemphylium blight severity data from the greenhouse in 2018 (*qSB-4-1*, yellow −LG4; *qSB-5-1* green – LG5), the growth chamber in 2019 (*qSB-2-1*, red-LG2) and in the field in 2019 (*qSB-2-2*, blue-LG2).

Four putative QTLs significantly associated with stemphylium blight resistance were identified in the LABC-01 population (Table 2, Figure 2). The individual QTLs explained 7.56% to 8.55% of the phenotypic variation. Two QTLs (*qSB-4-1* and *qSB-5-1*) were mapped on LG4 and LG5 based on mean severity scores from the greenhouse. The other two QTLs were mapped on LG2 based on mean severity scores from the growth chamber (*qSB-2-1*) and field (*qSB-2-2*), respectively. Three of these QTLs (*qSB-2-1, qSB-2-2*, and *qSB-4-1*) were inherited from the interspecific line LR-59-81, and one QTL, *qSB-5-1*, was inherited from the moderately resistant parent CDC Redberry. The flanking markers (1.5-LOD interval) of each QTL detected (Table 2, *qSB-2-1, qSB-4-1*, *qSB-5-1, qSB-5-2*) were used to anchor the four QTLs on the CDC Redberry reference genome. The physical QTL regions range from 0.13 Mbp (*qSB-2-1*) to 43.6 Mbp (*qSB-4-1*; Supplementary Table 2). The 0.13 Mbp region of QTL *qSB-2-1* contains seven genes, some of which were annotated as NBS-LRR proteins (*Lcu.2RBY.2g004820, Lcu.2RBY.2g004880*) and putative transmembrane proteins (*Lcu.2RBY.2g004830*). The region of the QTL *qSB-2-2* (10.40 Mbp) harbor 317 genes, some of which were described as Mitogen-activated protein kinase (Lcu.2RBY.2g009050), LRR receptor-like kinase (Lcu.2RBY.2g009850), Receptor Serine/Threonine kinase (*Lcu.2RBY.2g009530, Lcu.2RBY.2g008430*), Receptor-like kinase (*Lcu.2RBY.2g008700*), transmembrane protein (*Lcu.2RBY.2g008420*) and transcription factor (*Lcu.2RBY.2g009770*, *Lcu.2RBY.2g009750*). The 3.52 Mbp region of QTL *qSB-5-1* harbor 43 genes and the largest region 43.6 Mbp, of QTL *qSB-4-1* harbor 908 genes.

**Table 2.** Summary of quantitative trait loci (QTL) associated with stemphylium blight resistance identified using an advanced backcross population LABC-01 (n =182, BC_2_F_3:4_), derived from a cross between the *Lens culinaris* CDC Redberry × *L. ervoides* interspecific line LR-59-81. The population was screened under controlled environmental conditions in a greenhouse and a growth chamber, and under field conditions.

| Environment | QTL | LG<sup>a</sup> | Left Marker | Right Marker | 95%<br>Confidence<br>interval (cM)<sup>b</sup> | Peak<br>LOD<sup>c</sup> | %PVE<sup>d</sup> | ADD<sup>e</sup> |
| --- | --- | --- | --- | --- | --- | --- | --- | --- |
| Greenhouse-2018 | *qSB-4-1* | 4 | Lcu.2RBY.Chr4p376037412 | Lcu.2RBY.Chr4p419671210 | 520.2 - 524.5 | 4.04 | 8.55 | -0.53 |
|  | *qSB-5-1* | 5 | Lcu.2RBY.Chr5p314820775 | Lcu.2RBY.Chr5p311299110 | 25.5 - 40.5 | 3.65 | 7.56 | 0.29 |
| Growth chamber-2019 | *qSB-2-1* | 2 | Lcu.2RBY.Chr2p9874672 | Lcu.2RBY.Chr2p10001163 | 166.5 – 176.5 | 3.54 | 8.07 | -0.36 |
| Field-2019 | *qSB-2-2* | 2 | Lcu.2RBY.Chr2p16499628 | Lcu.2RBY.Chr2p26900380 | 181.5 - 190.5 | 3.28 | 8.30 | -2.97 |

<sup>a</sup> LG, Linkage group; <sup>b</sup> cM, centiMorgans; <sup>c</sup> LOD, logarithm of the odds value of peak marker; <sup>d</sup> %PVE, % of phenotypic variance explained by an individual QTL, <sup>e</sup> ADD, additive effects.

### 3.3 Population structure and genetic relatedness

Population structure was analyzed by DAPC and familial relatedness by pairwise kinship coefficient among the 101 *L. culinaris* accessions based on 143,971 genome-wide SNPs. These SNPs provide a whole genome-wide coverage along the 7 chromosomes of lentil cultivar CDC Redberry as shown in Figure 3. The SNP data was first transformed using a principal components analysis (PCA) and subsequently the number of clusters (sub-populations) was defined *a priori* according to three major climatic regions by DAPC (K = 3). Twenty PCs (58.4% of variance retained) of PCA and two discriminant eigenvalues were retained. These values were confirmed by cross-validation analysis. The biplot of PC1 by PC2 explaining 21.1% (PC1 = 13.2% and PC2 = 7.9%) of the total variance contained in the data shows overlaps of accession in the three major climatic regions (Figure 4). Accessions from the Mediterranean regions were the largest group (62), with some overlap with the temperate region group (25) and substantial overlap with the sub-tropical savannah group (14). The posterior probability plot confirmed this, with all three major climatic regions exhibiting admixtures (Figure 5). A phylogenetic tree and a heat map of the pairwise kinship values of the 101 accessions revealed a weak and complicated genetic relatedness among the genotypes (Figure 6). Both population structure and familial relatedness were included in the model of marker-trait association analysis to control inflation of p-values and false positive associations.

**Figure 3.**
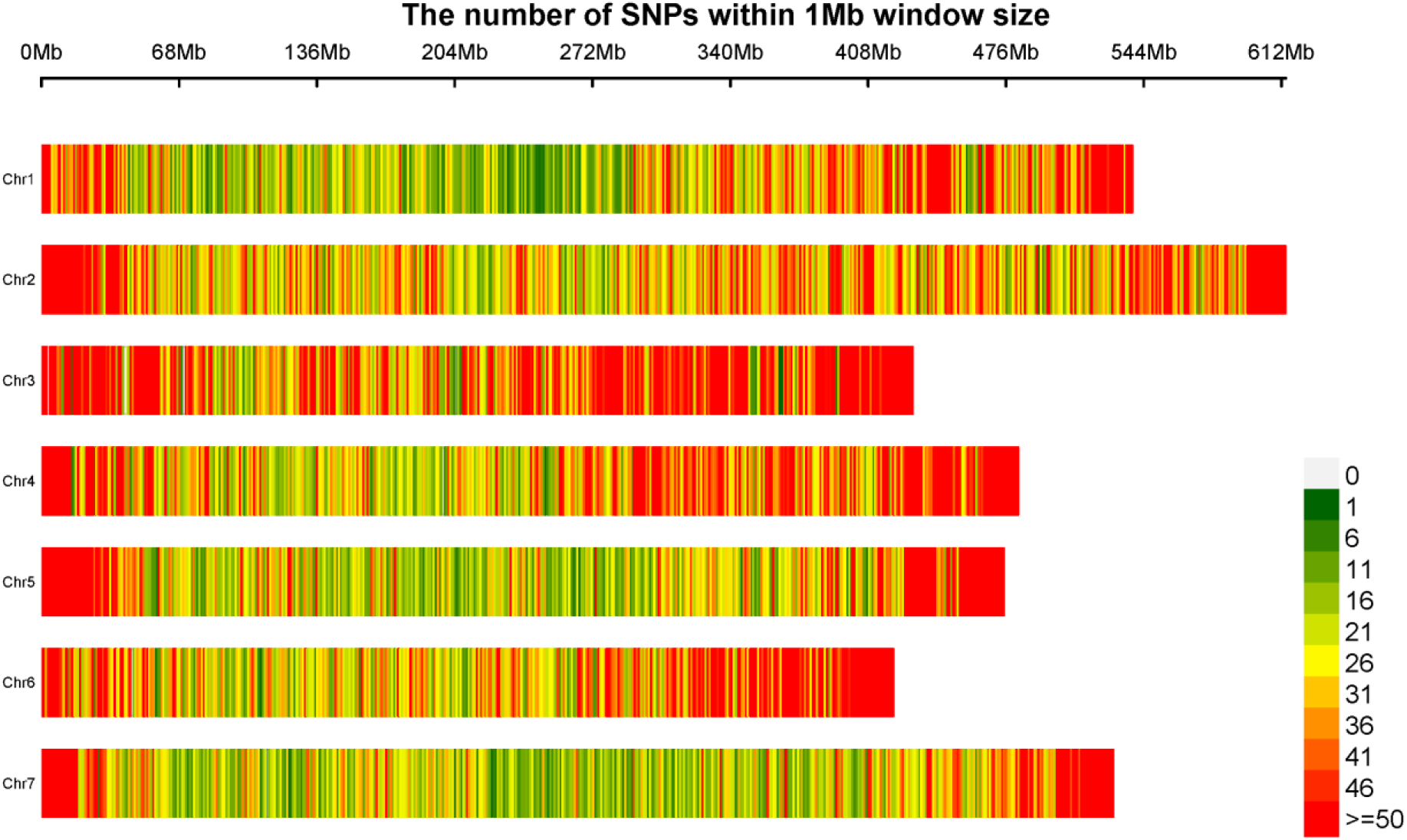
Plot of chromosome wise SNP density representing the number of SNPs within 1 Mb window size based on 143, 971 on the CDC Redberry genome v2.0. The horizontal axis indicates the chromosome length (Mb); the different colors depict SNP density.

**Figure 4.**
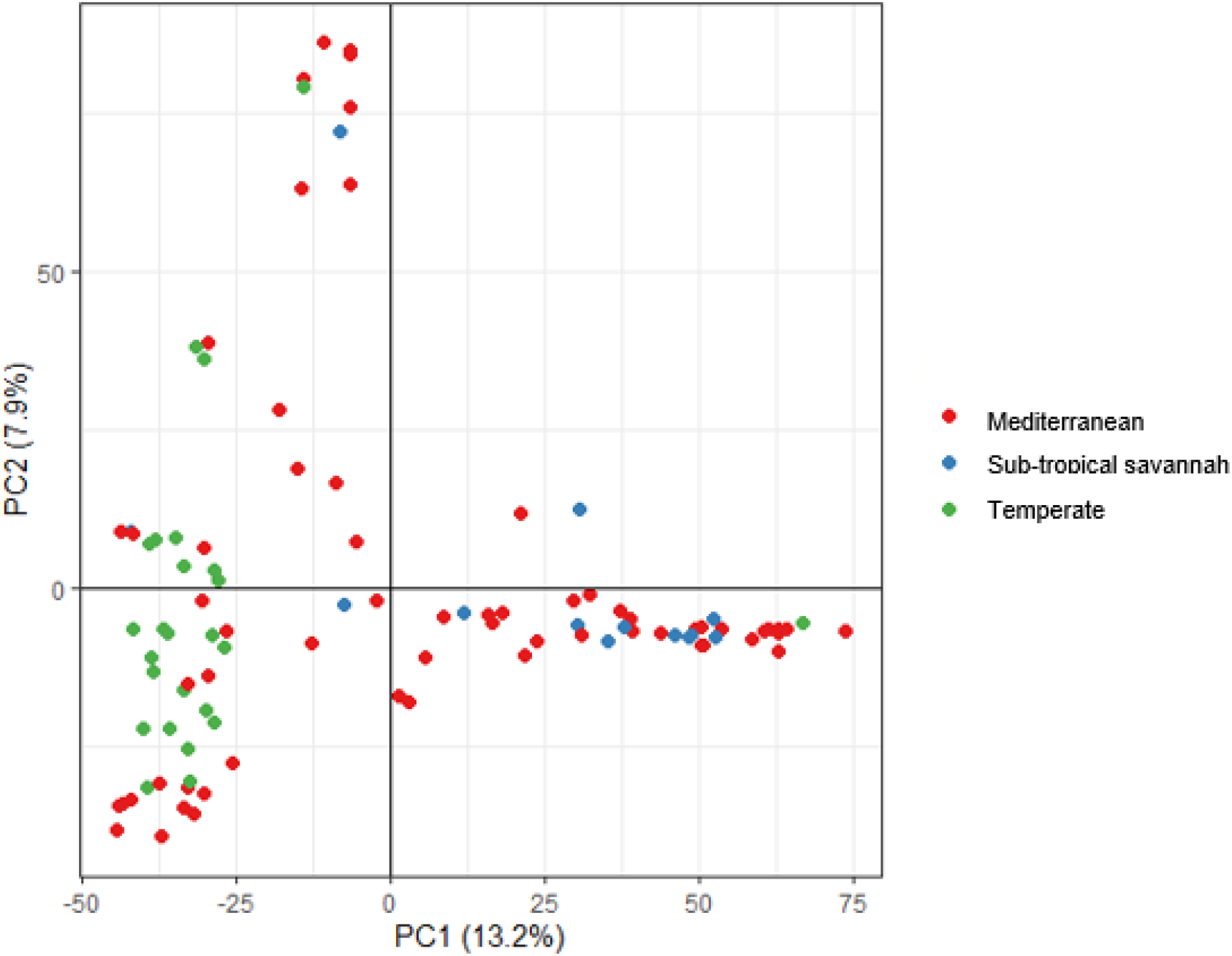
Plot of the first two principal components analyzed by discriminant analysis of principal components of the 101 lentil accessions based on 143,971 SNPs. Clusters were defined a priori by the three major climatic regions. The accession in the Mediterranean region forms a larger group with some mixed with the temperate region group and the rest mixed with the sub-tropical savannah group.

**Figure 5.**
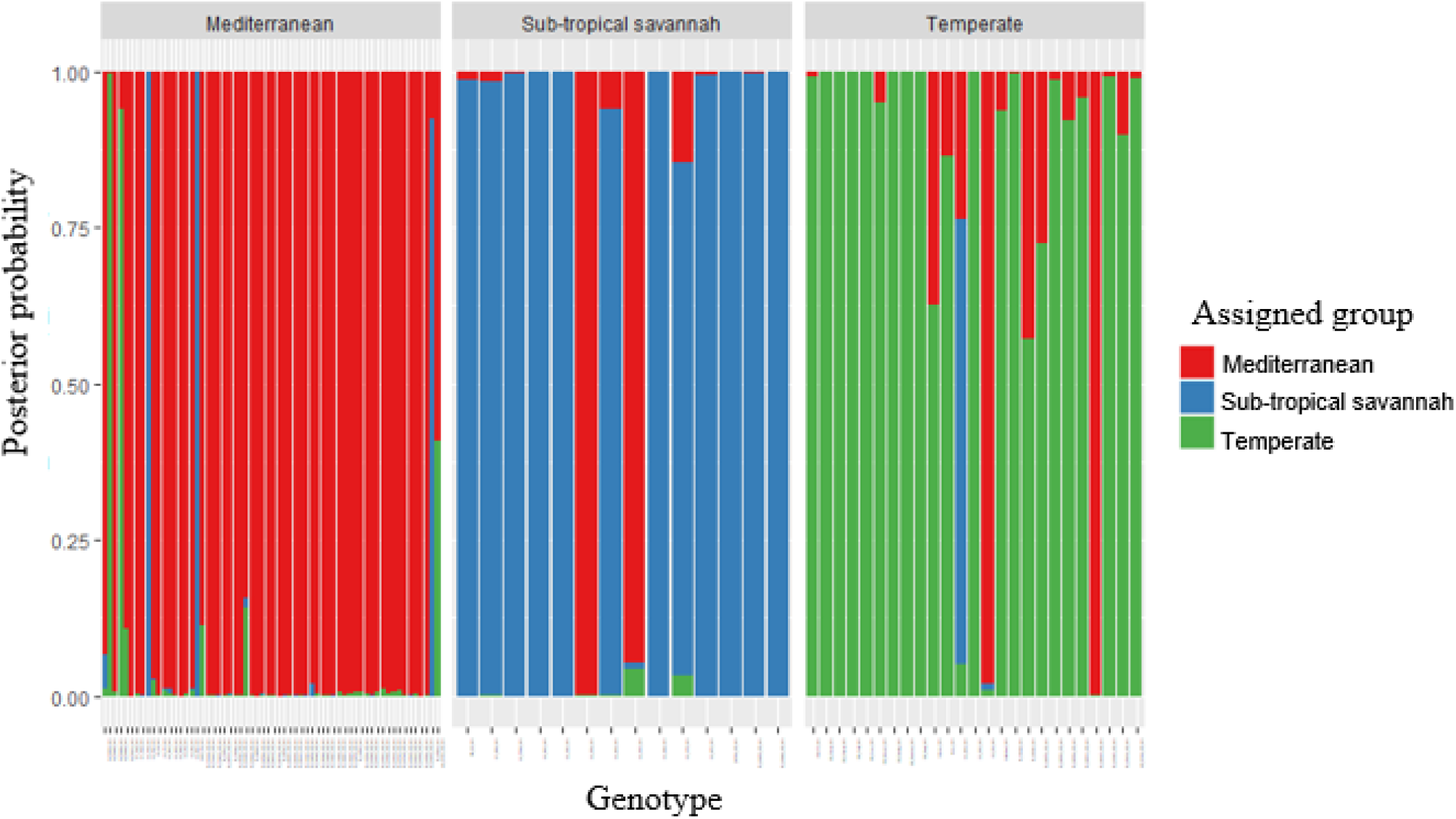
STRUCTURE-like plot of posterior probabilities for 101 lentil genotypes analyzed by discriminant principal component analysis, based on 143,971 genome-wide SNPs. A vertical colored line represents each individual, with same colored individuals belonging to same group. Population membership probability assignments by three major climatic regions (K = 3, Mediterranean, Sub-tropical savannah, and Temperate) exhibiting admixtures in all growing regions.

**Figure 6.**
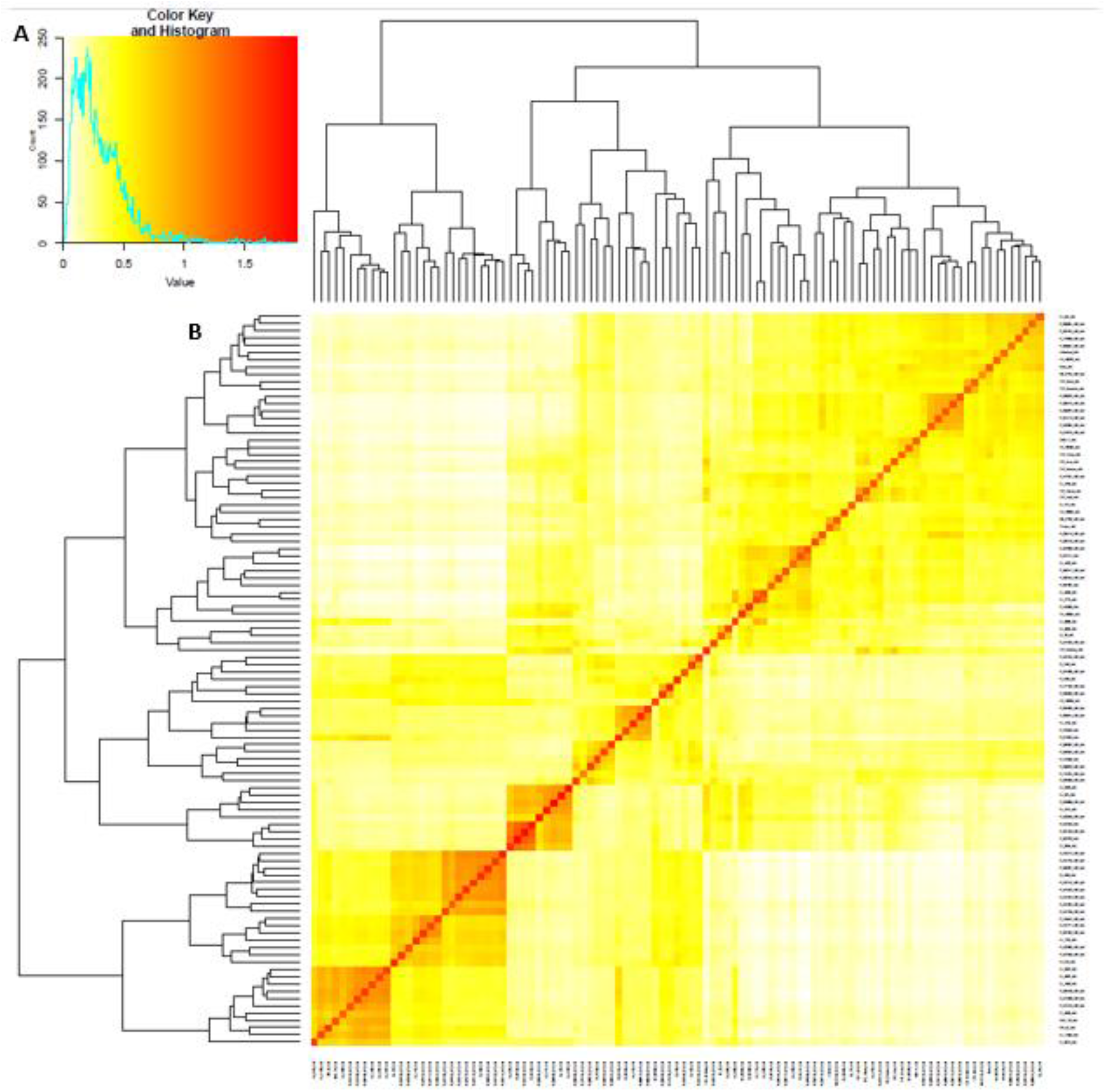
Plot of (A) histogram of kinship coefficient values and (B) hierarchical clustering and heat map of the pairwise kinship coefficient matrix values representing the level of genetic relatedness among 101 lentil accessions, based on 143, 971 SNP markers. The deep red boxes indicate highly related genotypes and orange to yellow indicate unrelated genotypes.

### 3.4 QTL-region-based marker-trait association analysis

We first conducted marker trait association analysis, focusing on the 2,416 SNP markers located in QTL *qSB-2-1, qSB-2-2, qSB-4-1, and qSB-5-1* identified in the bi-parental LABC-01 population (Table 2, Supplementary Table 2). The marker trait association was completed using FarmCPU model and the Q-Q-plot raised no concerns regarding genomic inflation, thereby confirming appropriate fitting of the model. Five SNP markers were detected to be significantly associated with stemphylium blight resistance (Table 3). The phenotypic variance explained by the five significant SNP markers ranged from 5.94% to 10.94%. One SNP marker was located within the QTL *qSB-2-1* region (Supplementary Figure 1), three within the QTL *qSB-4-1* region (Supplementary Figure 2), and one within the QTL *qSB-5-1* region (Supplementary Figure 3).

**Table 3.** Single nucleotide polymorphism markers associated with stemphylium blight resistance in QTL regions (QTL *qSB-2-1, qSB-4-1, qSB-5-1*) identified in advanced backcross population LABC-01 (n =182, BC_2_F_3:4_) using mean stemphylium blight severity means for a lentil diversity panel of 101 lentil accessions evaluated in the growth chamber. QTL-region marker-trait association analysis was performed with FarmCPU model.

| SNP Marker | Chromosome | Physical position (bp) | P-value | Minor allele frequency | R<sup>2</sup><sup>a</sup> |
| --- | --- | --- | --- | --- | --- |
| Lcu.2RBY.Chr2:9927887 | 2 | 9927887 | 7.57 x 10<sup>-06</sup> | 0.32 | 8.76 |
| Lcu.2RBY.Chr4:392720080 | 4 | 392720080 | 4.39 x 10<sup>-07</sup> | 0.19 | 10.94 |
| Lcu.2RBY.Chr4:413622606 | 4 | 413622606 | 5.31 x 10<sup>-08</sup> | 0.22 | 6.27 |
| Lcu.2RBY.Chr4:416496027 | 4 | 416496027 | 1.01 x 10<sup>-06</sup> | 0.27 | 5.94 |
| Lcu.2RBY.Chr5:313439017 | 5 | 313439017 | 5.67 x 10<sup>-07</sup> | 0.19 | 9.95 |

<sup>a</sup> % of phenotypic variance explained by each marker

Three SNP markers, Lcu.2RBY.Chr2:9927887, Lcu.2RBY.Chr4:392720080 and Lcu.2RBY.Chr4:416496027, were located within the gene models *Lcu.2RBY.2g004860* (Alpha-1, 4 glucan phosphorylase), *Lcu.2RBY.4g061330* (Auxin-induced protein 5NG4-like protein) and *Lcu.2RBY.4g065940* (Phosphoribosylformylglycinamidine cyclo-ligase) respectively, on chromosome 2 and 4 (Supplementary Table S3). LD analysis within 1 Mb of the significantly associated SNP marker revealed 27 SNP markers were in high LD (r^2^ > 0.7) with four out of the five SNPs (Supplementary Table 3). Sixteen of the 27 LD SNPs were located within six genes. Some of the genes were described as transcription factors (*Lcu.2RBY.5g043610*) and putative transmembrane proteins (*Lcu.2RBY.4g065310*).

### 3.5 Genome-wide association analysis

Genome-wide association analysis was performed on stemphylium blight severity scores and 143,971 SNP markers of the panel of 101 diverse lentil germplasms. The results of the GWAS showed that seven new significant SNP markers on chromosomes 1, 2, 3, 5, and 7 could be associated with stemphylium blight severity based on the FarmCPU model (Table 4, Figure 7). The phenotypic variance explained by the seven significant SNP markers ranged from 2.96% to 18.26%. LD analysis within 1 Mbp of the significantly associated SNP marker reveals 13 SNP markers had LD r^2^ > 0.7 (Supplementary Table 4). Four SNP markers, Lcu.2RBY.Chr3:44075187, Lcu.2RBY.Chr5:376522, Lcu.2RBY.Chr5:6575131 and Lcu.2RBY.Chr7:393853533, were located within the gene models *Lcu.2RBY.3g007400* (Acyl-CoA N-acyltransferase (NAT) family protein), *Lcu.2RBY.5g000250* (Neutral amino acid transporter), *Lcu.2RBY.5g004850* (Vacuolar protein sorting protein), and *Lcu.2RBY.7g050380* (LRR receptor-like kinase) respectively (Supplementary Table 4).

**Table 4.** Single nucleotide polymorphism markers associated with stemphylium blight resistance using mean stemphylium blight severity for a *Lens culinaris* diversity panel of 101 accessions evaluated in the growth chamber. Genome-wide marker-trait association analysis was performed with the FarmCPU model.

| SNP Marker | Chromosome | Physical position (bp) | P-value | Minor allele frequency | R<sup>2</sup><sup>a</sup> |
| --- | --- | --- | --- | --- | --- |
| Lcu.2RBY.Chr1:307105762 | 1 | 307105762 | 1.88 x 10<sup>-07</sup> | 0.12 | 3.37 |
| Lcu.2RBY.Chr2:4892081 | 2 | 4892081 | 6.22 x 10<sup>-08</sup> | 0.19 | 9.20 |
| Lcu.2RBY.Chr3:44075187 | 3 | 44075187 | 1.09 x 10<sup>-08</sup> | 0.31 | 8.03 |
| Lcu.2RBY.Chr5:376522 | 5 | 376522 | 3.34 x 10<sup>-10</sup> | 0.20 | 9.12 |
| Lcu.2RBY.Chr5:6575131 | 5 | 6575131 | 3.65 x 10<sup>-08</sup> | 0.07 | 11.87 |
| Lcu.2RBY.Chr5:393086848 | 5 | 393086848 | 2.40 x 10<sup>-10</sup> | 0.25 | 18.26 |
| Lcu.2RBY.Chr7:393853533 | 7 | 393853533 | 2.55 x 10<sup>-09</sup> | 0.08 | 2.96 |

<sup>a</sup> % of phenotypic variance explained by each marker

**Figure 7.**
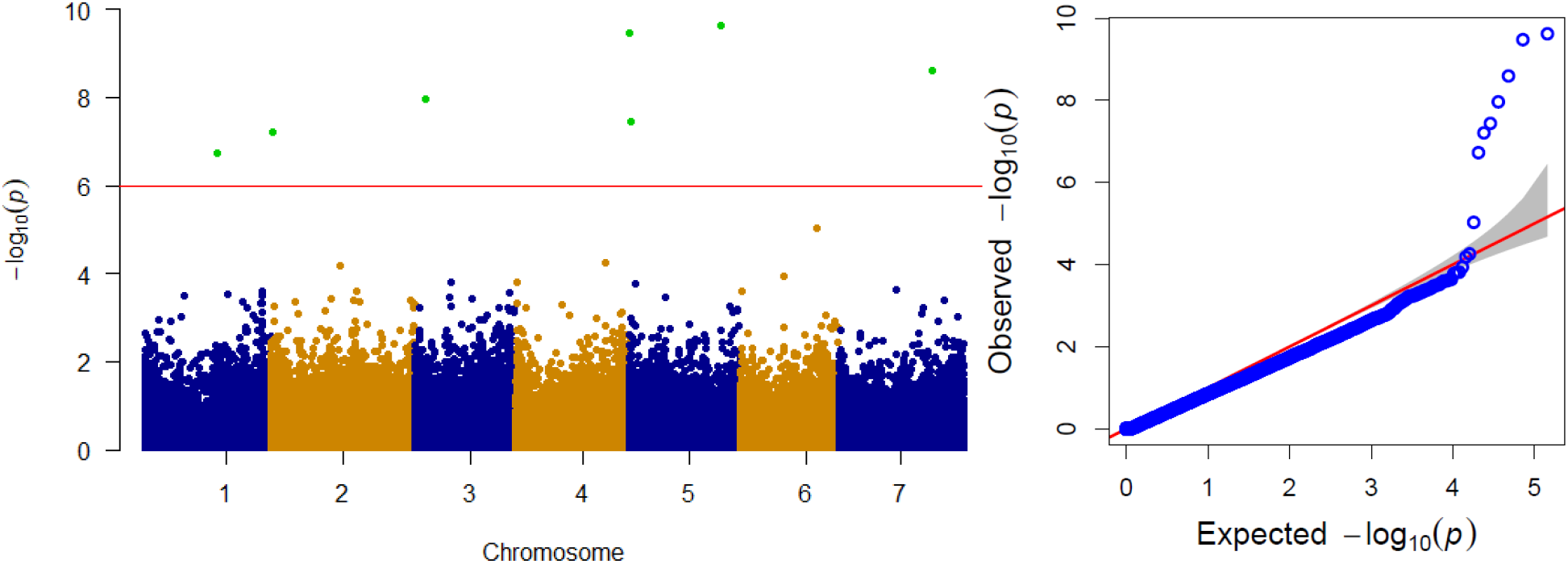
Manhattan plots showing seven SNP markers (green dots) associated with stemphylium blight severity based on FarmCPU model and Q-Q plots for test of association. Stemphylium blight severity data was collected from 101 *Lens culinaris* genotypes evaluated in the growth chamber. The red line indicates the significance threshold based on the Benjamini-Hochberg method for multiple test correction (FDR-adjusted *P* ≤ 0.05).

## 4 Discussion

Lentil is an important pulse crop with relatively low genetic diversity due to a genetic bottleneck created during domestication when it underwent selection for a small number of traits (Ladizinsky, 1987; Zohary, 1989). This has limited the genetic variation available in the cultivated gene pool for improving important agronomic traits. Several studies have reported *L. ervoides* accessions IG 72815 and L01-827A as a source of resistance to lentil diseases, including stemphylium blight (Tullu et al., 2010, 2013; Podder et al., 2013; Bhadauria et al., 2017; Adobor et al., 2020; Gela et al., 2021a). Introgression of genes for traits of interest from crop wild relatives or their interspecific hybrids into cultivated lentil through marker-assisted selection could help widen the genetic base and speed up the development of new cultivars with improved disease resistance. However, the transfer of stemphylium blight resistance from wild sources through marker-assisted selection requires deeper knowledge of its genetic basis and the development of markers linked to QTLs or genes regulating stemphylium blight resistance. In this research, linkage mapping was combined with LD mapping in the QTLs region and a genome-wide association study to identify QTLs and SNPs associated with resistance to stemphylium blight in lentil.

Although the distribution of stemphylium blight severity in the LABC-01 population indicated genotypes with severity scores lower than LR-59-81 and higher than CDC Redberry in all three environments, no significantly resistant transgressive segregants were observed in the population. The wide distribution of stemphylium blight severity suggests the population was suitable for QTL mapping even though both parents had some resistance to the disease, and that selection of genotypes with resistance alleles from both parents will benefit genetic improvement of stemphylium blight resistance in lentil. The frequency distribution of stemphylium blight severity in the LABC-01 population showed a unimodal distribution, indicating a polygenic inheritance of stemphylium blight resistance. Similarly, Bhadauria et al. (2017) reported polygenic inheritances of stemphylium blight resistance in the intraspecific *L. ervoides* RIL population LR-66 derived from the cross IG 72817 × L01-827A. Research on a *L. culinaris* RIL population derived from the cross ILL 6002 × ILL 5888 screened in the field in Bangladesh indicated that stemphylium blight resistance was controlled by dominant genes (Saha et al., 2010). The differences might be due to the different genetic backgrounds involved, environmental effects or the different isolates of *Stemphylium spp* infecting lentil in the field in Bangladesh, and/or the different disease severity rating scale.

The proportion of markers introgressed from the donor genotype (9.5%) was almost equal to the expected amount of 12.5% for a BC_2_ population in the absence of selection (Stam and Zeven, 1981). Four putative QTLs regulating stemphylium blight resistance in the LABC-01 population were identified: two on chromosome 2, one on chromosome 4, and one on chromosome 5 of the *L. culinaris* genome. The QTLs detected could be classified as minor effect QTLs with phenotypic variation ranging from 7.56% to 8.55%, underlining the complex polygenic inheritance of the trait. Furthermore, the four QTLs for stemphylium blight resistance identified in this study were environment-specific. Saha et al. (2010) also reported a strong environmental influence on stemphylium blight resistance, identifying one QTL based on stemphylium blight severity score in a *L. culinaris* RIL population in 2007 and three QTLs for the same population in 2009. This was further confirmed in the interspecific *L*. *culinaris × L. ervoides* RIL population LR-26 by Adobor et al. (2020). QTLs consistently detected across different environments are ideal for introgression and use in breeding programs because crops with such QTLs are more likely to display a resistant phenotype in more than one environment (El-Soda et al., 2014). However, environment-specific QTLs can also be useful since breeding programs do not cover all regions where the crop is grown but are focused on regions with similar environments (El-Soda et al., 2014).

Validation of QTLs across different populations is important because of differences in allele effects depending on the genetic background (Tanksley and Hewitt, 1988). A previous QTL analysis identified three significant QTLs (two on chromosome 2 and one on chromosome 3) linked with stemphylium blight resistance in intraspecific *L. ervoides* population LR-66 (Bhadauria et al., 2017). Due to the unavailability of an *L. ervoides* genome at the time, SNP markers of that study were identified based on an earlier version of the *L. culinaris* CDC Redberry genome. The limitations imposed by the use of different CDC Redberry genome versions for SNP marker identification and QTL analysis preclude the direct comparison to determine if QTLs detected on chromosome 2 in both studies co-localize in the same region. The detection of QTLs on chromosomes 4 and 5 and the absence of QTLs on chromosome 3 in the current study suggests that some of the QTLs were probably specific to the population and the parents involved (Han et al., 2012). Thus, the introgression or pyramiding of these QTLs from different resistance sources into one genetic background is important to improving resistance.

LD mapping can be used to systematically scan for candidate loci for association with a trait of interest across the whole genome or in an interval (region) defined by linkage mapping. A limitation of GWAS is that SNP markers that may be associated with the trait do not reach statistical significance because of their small effect sizes and thus may not be detected. In this study, marker trait association analysis was performed on four QTL regions (*qSB-2-1, qSB-2-2, qSB-4-1*, and *qSB-5-1*) defined by markers identified from linkage mapping in the LABC-01 population. Five significant SNP markers were identified that were associated with stemphylium blight severity on chromosomes 2, 4, and 5. In addition, the genome-wide association analysis identified an additional seven significant SNP markers on chromosomes 1, 2, 3, 5, and 7 outside the QTL regions on the linkage map of LABC-01.

The GWAS approach is suited for refining the genomic regions with high resolution as it captures a larger portion of the recombination events that have accumulated in the diversity panel (Zhu et al., 2008). Thus, the SNPs uncovered by using both methods will facilitate pyramiding of the resistance genes from diverse sources into lentil cultivars to achieve high levels of stemphylium blight resistance. Some of the SNPs in high LD with significant SNP markers were located within genes described as LRR receptor-like kinase (*Lcu.2RBY.7g050380*), putative transmembrane protein (*Lcu.2RBY.4g065310*) and transcription factor (*Lcu.2RBY.5g043610*). These genes might play an important role in gene transcription, signaling and defense response to pathogens (Afzal et al., 2008; Thomma et al., 2011; Hamel et al., 2012; Jiang et al., 2018; Campos et al., 2022). The LD SNP markers located within these genes can be candidate markers for validation.

Accurate phenotyping is one of the major constraints in genetic studies of disease resistance and resistance breeding, especially in semi-arid areas of Saskatchewan and Alberta, the prairie provinces of Western Canada where about 40% of the world’s lentil is produced. Evaluation of disease resistance on lentil plants grown in small pots under controlled conditions allows for screening of a large number of genotypes. Disease severity assessments in a growth chamber or greenhouse may reflect the genotypic effect on stemphylium blight more so than any environmental effects. However, the artificial conditions in a growth chamber or greenhouse do not accurately reflect field conditions; therefore, results need to be validated in the field. In this research, attempts were made to evaluate the lentil diversity panel in the field, but high temperatures and dry conditions in the 2020 and 2021 growing seasons limited plant infection and disease development even with the use of artificial inoculation, covering of plots with low-tunnel plastics, and application of sprinkler irrigation.

To conclude, we combined linkage mapping and genome-wide marker trait association analysis to dissect genetic control of stemphylium blight resistance in populations originating from *L. ervoides* interspecific and *L. culinaris*. Four QTLs controlling stemphylium blight resistance were detected in the bi-parental LABC-01 population evaluated in three environments. The resistant LABC-01 genotypes are useful sources of resistance for the breeding of stemphylium blight resistant cultivars because they carry QTL alleles identified in the adapted background. Furthermore, twelve SNP markers were identified to be associated with stemphylium blight resistance in a lentil diversity panel. The significant SNP markers associated with stemphylium blight resistance can be useful for predictive phenotypic selection of stemphylium blight resistant genotypes through marker-assisted selection after validation.

## 5 Acknowledgements

We acknowledged funding from the Saskatchewan Pulse Growers and the Agricultural Development Fund of the Saskatchewan Ministry of Agriculture. Special thank you to Brent Barlow and Kevin Mikituk for technical assistance in the field. We appreciate the Pulse Crop Genetics and Breeding lab members Akiko Tomita and Robert Stonehouse for their assistance with exome capture library preparation for sequencing. We thank Larrisa Ramsay and Carolyn Caron for assistance on bioinformatics. We are also grateful for genetic resources made available from the ‘Application of Genomics to Innovation in Lentil Economy (AGILE)’ project.

