## Supplementary Tables for "Mapping of genomic regions linked to stemphylium blight (*Stemphylium botryosum* Wallr.) resistance in lentil using linkage mapping and marker-trait association analysis"

### 1 Supplementary figures

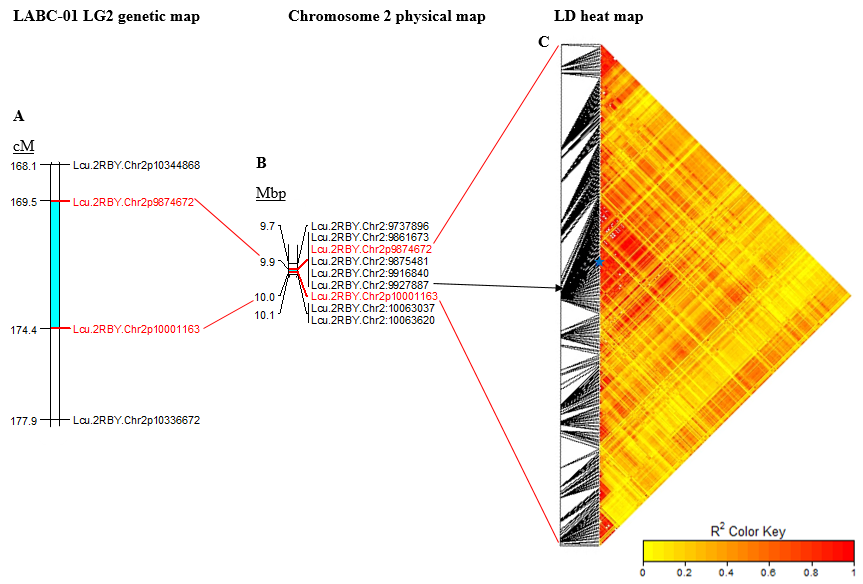

Supplementary Figure 1. Position of the stemphylium blight resistance QTL *qSB-2-1* on (A) linkage group 2 genetic map of the advanced backcross population LABC-01, evaluated in the growth chamber and on (B) physical map region (0.13 Mbp) on chromosome 2 of CDC Redberry genome v2.0. The positions are in centiMorgans (cM) and mega base pairs (Mbp) as indicated at the top of the bar. (C) Heat map plot of marker pairs linkage disequilibrium (LD) within a 1 Mbp window of one SNP marker (black arrows) associated with stemphylium blight resistance in the QTL *qSB-2-1* region (B) containing 188 SNP markers. The blue dot is the starting and ending points for SNP associated with stemphylium blight resistance. The colored fill represents marker pairwise LD measure as r^2^. The darkest color cells are indicative of the highest LD.

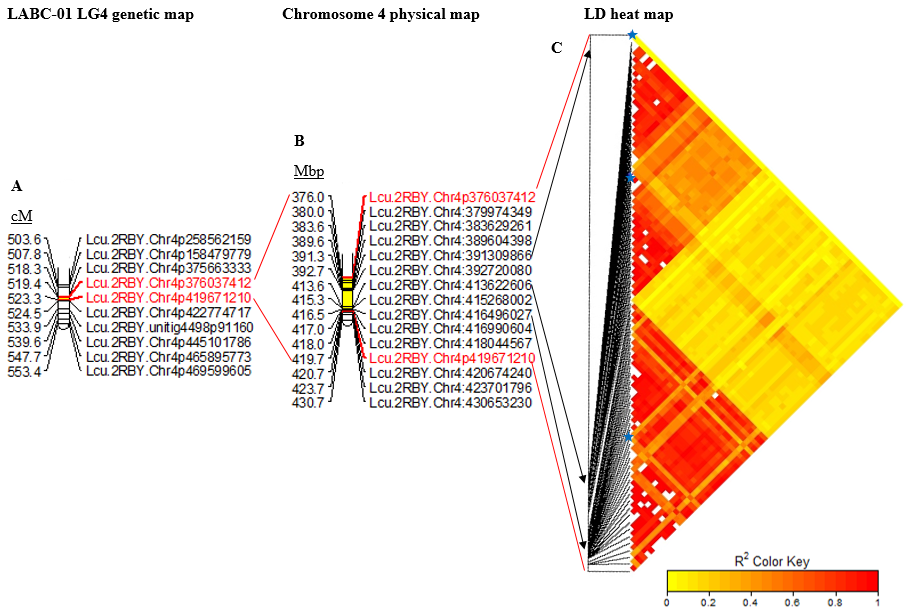

Supplementary Figure 2. Position of the stemphylium blight resistance QTL *qSB-4-1* on (A) linkage group 4 genetic map of the advanced backcross population LABC-01 evaluated in the greenhouse and on (B) physical map region (43.6 Mbp) on chromosome 4 of CDC Redberry genome v2.0. The positions are in centiMorgans (cM) and mega base pairs (Mbp), as indicated at the top of the bar. (C) Heat map plot of pairwise marker linkage disequilibrium (LD) within a 1 Mbp window of three SNP markers (black arrows) associated with SB resistance in the QTL *qSB-4-1* genomic region (B) containing 67 SNP markers. The blue dots are the starting and ending points for SNP with associated stemphylium blight resistance. The colored fill represents marker pairwise LD measure as r^2^. The darkest color cells are indicative of the highest LD.

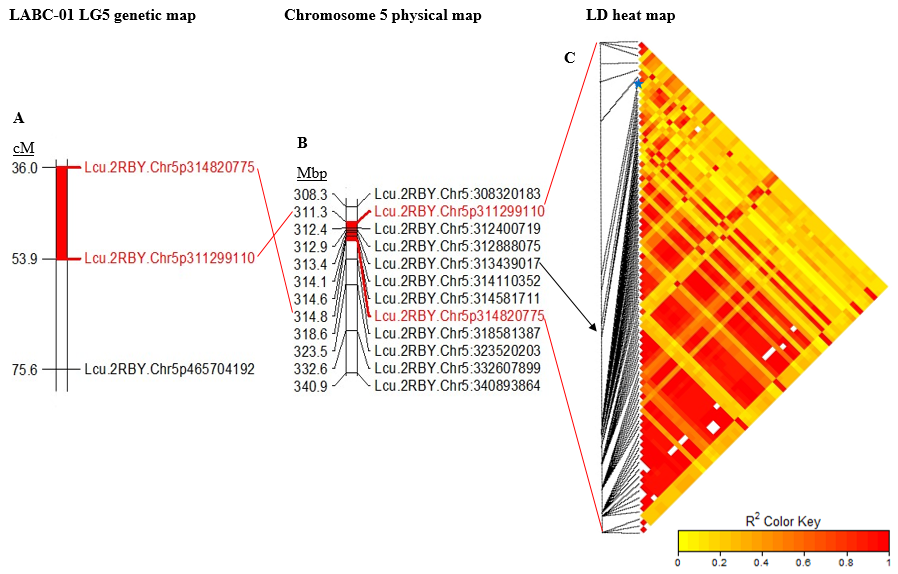

Supplementary Figure 3. Position of stemphylium blight resistance QTL *qSB-5-1* on (A) linkage group 5 genetic map of the advance backcross population LABC-01 evaluated in the greenhouse and on (B) physical map region (3.6 Mbp) on chromosome 5 of CDC Redberry genome v2.0. The positions are in centiMorgans (cM) and mega base pairs (Mbp) as indicated at the top of the bar. (C) Heat map plot of marker pairs linkage disequilibrium (LD) within a 1 Mbp window of SNP marker (black arrow) associated with stemphylium blight resistance in the QTL *qSB-5-1* region on chromosome 5 containing 71 SNP markers. The blue dot is the starting and ending points for SNP with associated stemphylium blight resistance. The colored fill represents marker pairwise LD measure as r^2^. The red color cells are indicative of the highest LD.
